# Activating M1 muscarinic cholinergic receptors induces destabilization of resistant contextual fear memories in rats

**DOI:** 10.1101/2023.05.18.541305

**Authors:** Karim H. Abouelnaga, Andrew E. Huff, William S. Messer, Boyer D. Winters

## Abstract

Destabilization of previously consolidated memories places them in a labile state in which they are open to modification. However, strongly encoded fear memories tend to be destabilization-resistant and the conditions required to destabilize such memories remain poorly understood. Our lab has previously shown that exposure to salient novel contextual cues during memory reactivation can destabilize strongly encoded object location memories and that activity at muscarinic cholinergic receptors is critical for this effect. In the current study, we similarly targeted destabilization-resistant fear memories, hypothesizing that exposure to salient novelty at the time of reactivation would induce destabilization of strongly encoded fear memories in a muscarinic receptor-dependent manner. First, we show that contextual fear memories induced by 3 context-shock pairings readily destabilize upon memory reactivation, and that this destabilization is blocked by systemic (ip) administration of the muscarinic receptor antagonist scopolamine (0.3mg/kg) in male rats. Next, we demonstrate that more strongly encoded fear memories (induced with 5 context-shock pairings) resist destabilization. Consistent with our previous work, however, we report that salient novelty (a change in floor texture) presented during the reactivation session promotes destabilization of resistant contextual fear memories in a muscarinic receptor-dependent manner. Finally, the effect of salient novelty on memory destabilization was mimicked by stimulating muscarinic receptors with the selective M1 agonist CDD-0102A (ip, 0.3mg/kg). These findings reveal further generalizability of our previous results implicating novel cues and M1 muscarinic signaling in promoting destabilization of resistant memories and suggest possible therapeutic options for disorders characterized by persistent, maladaptive fear memories such as PTSD and phobias.

## 1. Introduction

The retention of newly acquired information in long-term memory requires a protein synthesis-dependent process to transform neural representations from a labile state to a stable, consolidated memory trace (McGaugh, 2000; Nader, 2015; Nader, 2000). When memories are recalled or reactivated, they can return to a destabilized state in which they are open to modification through strengthening, weakening, or other forms of information integration (Nader, 2000; Nader, 2015). Following this, further synaptic changes can restabilize the updated memory trace in a process known as reconsolidation (Li et al, 2013).

Maladaptive memories such as post-truamtic stress disorder (PTSD) and phobias caused by dysregulated fear circuits (Grella et al., 2022; Chen et al., 2021) could be the result of a breakdown of reactivation-induced memory updating (Feduccia and Mithoefer, 2018; Milad et al., 2008). In rats, strongly encoded fear memories tend to resist destabilization when a simple reactivation cue is presented (Suzuki et al., 2004; Wang et al., 2009). Additionally, resistance to destabilization has been observed for other types of memories such as strongly encoded object location or recognition memories (Huff et al., 2022; Winters et al., 2009) or relatively remote memories (Eisenberg & Dudai, 2004; Stiver et al., 2015; Wideman et al., 2021). This indicates that when a memory is strongly encoded or less recently acquired, it is subject to boundary conditions which render the simple reminder-cue reactivation protocol insufficient to induce destabilization (Winters et al., 2009; Bui and Milton, 2023).

However, such boundary conditions are not immutable. Consistent with the notion that memory reconsolidation serves an updating function (Lee, 2009; Lee et al., 2017; Schiller et al., 2010), past studies have indicated that prediction error or salient novel cues presented at the time of memory reactivation can render otherwise resistant memories labile (Sinclair and Barense, 2018, 2019; Sinclair et al., 2021; Winters et al, 2009; Morris and Jones, 1990; Fernández and Morris, 2018). Indeed, in our hands, the introduction of salient novelty during reactivation induces the destabilization of both resistant object recognition memories in perirhinal cortex (PRh) and resistant object location memories in the dorsal hippocampus (dHPC; Stiver et al, 2015, 2017; Huff et al, 2022). By extension, the addition of novel salient information during reactivation could be a possible method to induce destabilization of resistant fear memories. Accordingly, strongly encoded fear memories can be destabilized by increasing the length of the reactivation session (Suzuki et al., 2004) or introducing temporal novelty (Diaz-Mataix et al., 2013), but the variety of stimuli that can induce such effects, and their neurobiological bases, have not been thoroughly explored.

We have previously implicated the neurotransmitter acetylcholine in the destabilization of strongly encoded object location and remote object recognition memories (Stiver et al, 2015, Huff et al, 2022, Winters et al, 2009). Acetylcholine has an established role in multiple cognitive processes related to learning, such as arousal, attention, encoding, and novelty detection (Sarter and Bruno, 1997; Hasselmo and McGaughy, 2004; Sturgill et al., 2020), and cholinergic transmission could therefore be promoted by the kinds of stimuli that induce memory destabilization. Indeed, scopolamine, a muscarinic acetylcholine receptor (mAChR) antagonist, blocks novelty-induced destabilization of object memories when infused into PRh (Stiver et al, 2015), and object location memories when infused into the dHPC (Huff et al., 2022). Moreover, infusion of CDD-0102A, an M1 specific mAChR agonist, into the dHPC or PRh prior to reactivation promotes destabilization of strongly encoded object location memories (Huff et al, 2022) and remote object memories (Stiver et al., 2017), respectively. Thus, the cholinergic system plays a critical role in the destabilization of object-based memories that depend on either PRh or the dHPC. However, the involvement of the cholinergic system in destabilization of fear memories has not been explored.

Here, we used contextual fear conditioning (CFC) to investigate the role of the cholinergic system in destabilization of strongly encoded fear memories. Specifically, we aimed to uncover the role of mAChRs in promoting the destabilization of strong fear memories. While there is evidence to support the role of mAChRs in fear memory reconsolidation (Krawczyk et al., 2021), to our knowledge, their role in destabilization has not been explored. We hypothesized that - similar to their role in the destabilization of object and object location memories – mAChR activation is necessary for fear memory destabilization; thus, systemic administration of scopolamine or CDD-0102A should block or promote fear memory destabilization, respectively. Additionally, we predicted that salient novelty would induce destabilization of strongly encoded contextual fear memory and that this effect should also be mAChR-dependent.

## 2. Methods

### 2.1. Subjects

184 male Long-Evans rats were obtained from Charles River, QC and arrived weighing between 150g and 250g. Testing began when rats reached an approximate weight of 275g. Upon arrival, they were placed in standard rectangular cages (48 x 26 x 20cm) composed of polycarbonate and containing Envirodry bedding and standard enrichment (nesting materials and wooden block). They were housed with a reversed light/dark cycle in which lights turned off at 8:00 and turned on at 20:00. Rats were acclimated to the room for one week before experiencing researcher handling in preparation for the experiments. Food and water were available *ad libitum*. Behavioral testing took place during the dark phase. All procedures were approved by the Animal Care Committee (ACC) of the University of Guelph.

### 2.2. Drug Administration

The non-selective NMDA receptor antagonist, MK-801 (Sigma Aldrich, Ontario, Canada), was dissolved in 0.9% physiological saline and delivered at a dosage of 0.2 mg/kg intraperitoneally (ip). When administered systemically, this drug blocks NMDA receptors which prevents the process of reconsolidation, creating a memory impairing effect (Wideman et al., 2022; Winters et al., 2009; Stiver et al., 2015; Benvenga and Spaulding, 1988; Brown et al., 2008; Zaichenko et al., 2021; Przybyslawski & Sara, 1997). Recent evidence indicates that NMDA receptors contribute to both the process of destabilization and reconsolidation (Wideman et al., 2020; Milton et al., 2013). To limit this drug from affecting destabilization, it was administered systemically immediately prior to reactivation to ensure that its peak activation would occur during the reconsolidation window following memory reactivation (Wideman et al., 2022).

The non-selective muscarinic receptor antagonist scopolamine hydrobromide (SCOP; Sigma Aldrich, Ontario, Canada) was dissolved in 0.9% physiological saline and given ip at 0.3 mg/kg based on past evidence that scopolamine successfully blocks destabilization at this dose (Stiver et al., 2015; Wideman et al., 2022; Flavell et al., 2013).

To promote destabilization, the highly selective M1 muscarinic receptor agonist CDD-0102A (generously provided by Dr. William Messer, Jr., University of Toledo, Toledo, OH) was dissolved in 0.9% physiological saline and administered at a dose of 0.3mg/kg, ip. CDD-0102A was previously utilized in promoting destabilization of object memories when administered systemically in mice (Jardine et al., 2022) and intracranially in rats (Wideman et al., 2023). Physiological saline (0.9%) was used as a control (VEH) in each experiment.

### 2.3. Fear Conditioning Apparatus

Four fear conditioning boxes (30cm x 24cm x 24cm), each housed within separate sound-attenuating chambers (Med Associates Inc., Fairfax, VT), were used for all experiments. Near infrared imaging (NIR) was accomplished via a camera mounted inside the door of each chamber. The camera captured movement at 30 frames per second (fps) and automatically scored freezing through Video Freeze software (Med Associates Inc., Fairfax, VT). Throughout the experiments, the house light was always on in the boxes. Between rats, boxes were cleaned with 5 % hydrogen peroxide (H_2_O_2_).

### 2.4. Standard Fear Conditioning Protocol

The standard fear conditioning protocol consisted of three sessions that were 24 h apart. Before starting the sessions, rats were habituated to injections and transport for a period of two days. The training session took place 24 h following the second habituation day. For this session, the rats were placed in the chamber for a two min baseline period during which no foot-shocks occurred. Following this, three foot-shocks (1 mA, 1s each) were delivered 60s apart. This was followed by a 2-min post-shock period during which no foot-shock was presented before the rat was removed from the box. The total training session time was 6 minutes.

The next day, the contextual fear memory was reactivated for each rat in its original training box. Prior to reactivation, the rats received injections based on their assigned group (see specific details below). After this, they were placed in the chamber for a 90-s period to reactivate the memory. There were no foot-shocks delivered in this session. The test session occurred 24h later when the rats were placed back in the fear conditioning box for an 8-minute session. There were no foot-shocks presented, and all freezing behavior was automatically scored through the behavioral recording software Video Freeze.

#### 2.4.1 Resistant Fear Conditioning Protocol

To establish a strongly encoded fear memory, the standard fear conditioning protocol was modified. The number of foot-shocks given on the training session increased from three to five. Therefore, the training session was extended from six minutes to eight minutes in total. For the reactivation and test sessions, all parameters remained the same.

### 2.5 Experimental Approach

#### 2.5.1 Experiment 1

The purpose of this experiment was to determine whether the contextual fear memory established by the standard fear conditioning protocol readily destabilizes when reactivated. To establish that, the NMDA receptor antagonist, MK-801 was used to inhibit reconsolidation by injecting it immediately prior to reactivation (Winters et al., 2009; Wideman et al., 2022; Lee et al., 2006; Stiver et al., 2015). A group of 24 rats received either pre-reactivation MK-801, pre-reactivation VEH or MK-801 with no reactivation session (n = 8 per group).

#### 2.5.2 Experiment 2

There is past evidence that muscarinic cholinergic receptors (mAChR) play a key role in blocking the destabilization of memories in object-based tasks (Huff et al., 2022, Wideman et al., 2022, Stiver et al., 2015). Experiment 2 aimed to determine whether this also applies for contextual fear conditioning. This experiment involved administering scopolamine 20 min prior to reactivation in conjunction with either MK-801 or VEH immediately prior to reactivation. Scopolamine was administered 20 min earlier to allow it to cross the blood-brain barrier and block the destabilization of contextual fear memory. This experiment incorporated 32 rats divided into four experimental groups (SCOP-MK-801, SCOP-VEH, VEH-MK-801, VEH-VEH). The drug denoted first was administered 20 min prior, while the second listed drug was administered immediately prior to reactivation.

#### 2.5.3 Experiment 3

The purpose of this experiment was to create a contextual fear memory that is resistant to destabilization. A new cohort of 32 rats was divided equally into groups receiving either 3 or 5 foot-shocks. Half the rats received VEH and half were given MK-801 immediately prior to reactivation (n = 8 per group).

#### 2.5.5 Experiment 4

Following the third experiment, the remaining experiments used the resistant fear conditioning protocol. For Experiment 4, the aim was to promote the destabilization of a strongly encoded contextual fear memory. Past evidence has indicated that the presentation of novelty during memory reactivation in the form of a floor insert promotes the destabilization of spatial and object memories (Huff et al., 2022, Stiver et al., 2015; Winters et al., 2009). Accordingly, for this experiment we placed a novel floor insert inside the chamber during memory reactivation for half the rats. The insert changed the tactile nature of the metal circular rods in the fear conditioning chamber to a uniform floor made of white plastic. The floor insert covered the entirety of the floor of the chamber. A new cohort of 32 rats was equally divided into groups either receiving novelty during reactivation (NOV) or not. Furthermore, within each group, half the rats received VEH and half MK-801 immediately prior to reactivation.

#### 2.5.6 Experiment 5

The purpose of this experiment was to determine whether mAChRs play a role in novelty-induced destabilization of resistant contextual fear memories. For this experiment, a new cohort of 32 rats was used. All rats were exposed to a novel floor insert that was added during the reactivation session. Twenty minutes prior to reactivation, two groups received a scopolamine (SCOP) injection and 2 other groups received VEH; immediately prior to reactivation, the 4 groups received either MK-801 (SCOP-MK-801, VEH-MK-801) or VEH (SCOP-VEH, VEH-VEH).

#### 2.5.7 Experiment 6

For this last experiment, the aim was to promote the destabilization of resistant contextual fear memory by activating M1 mAChRs systemically. Past evidence has implicated M1 mAChR agonism in the destabilization of strongly encoded object location memories (Huff et al., 2022), as well as remote object memories (Stiver et al., 2017), in the absence of explicit contextual novelty during memory reactivation. For this reason, the M1 mAChR agonist CDD-0102A was used to promote destabilization. Thirty-two naïve rats were used for this experiment. Sixteen rats received ip CDD-0102A injections 30 minutes prior to reactivation to ensure it crossed the blood-brain barrier. The remaining 16 rats received VEH injections. Immediately prior to reactivation, the rats received either MK-801 or VEH. For this experiment, there was no explicit novelty included during memory reactivation.

### 2.6 Data Analysis

All freezing behavior was collected using Med Associates Inc’s stock software, Video Freeze. Freezing was defined as no movement except for breathing (Maren, 2001). Freezing was automatically scored by the software in real time and verified by the researcher for discrepancies. Raw data files were analyzed using Video Freeze’s internal component analysis function, which bases each analysis on the session’s duration and parameters. Freezing behavior was denoted as precent freezing out of 100% based on the timing of each entire session. One-way analysis of variance (ANOVA) was utilized to determine main group effects for experiment one. For experiments 2-6, 2×2 ANOVAs were used. Tukey post-hoc tests were conducted for multiple comparisons. In addition, independent samples t-test were used for experiments with two groups. All statistical analyses were conducted with IBM SPSS (Version 28).

## 3. Results

### 3.1. Contextual fear memories are readily destabilized when briefly reactivated

The standard fear conditioning protocol was used in which rats were trained with three shocks and the memory was briefly reactivated 24h later. MK-801 reduced freezing behavior in the test session when given immediately prior to reactivation (Fig. 1A). Results of a one-way between-subjects ANOVA indicated a significant main effect of drug (*F* (2,20) = 7.920, *p* = 0.0029, η^2^ = 0.44), and Tukey’s multiple comparisons test revealed that MK-801 significantly reduced freezing in the test session compared to the VEH (*p* = 0.0068) and the No-RA (*p* = 0.0076) groups, which did not differ (*p* = 0.9855).

**Fig. 1.**
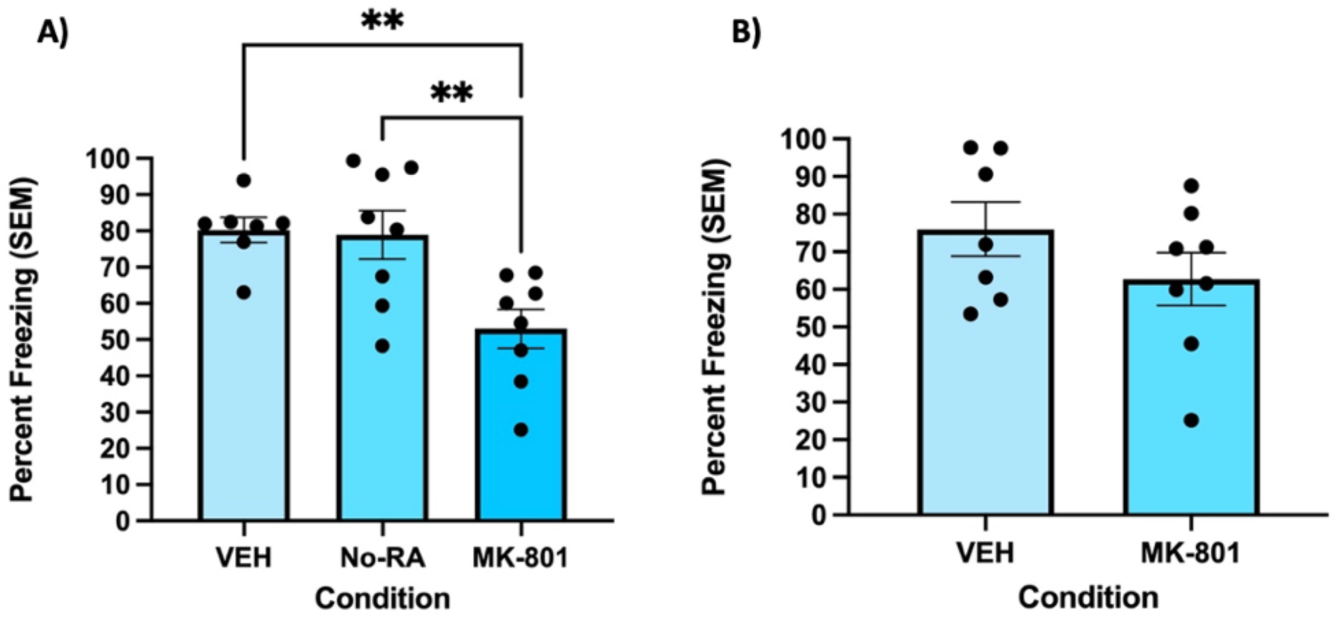
Contextual fear memories are destabilized when reactivated, and MK-801 blocks the reconsolidation of the memory in a reactivation dependent manner. A) In the test session, rats that received MK-801 prior to memory reactivation displayed significantly less freezing compared to the VEH and No-RA groups. Freezing behavior (n = 8 per group) was measured as percent freezing (± SEM) across an 8-min test session. B) During the reactivation session, there was no significant difference between the VEH and MK-801 groups suggesting no significant effect of drug on the expression of freezing, and that rats successfully learned the context – shock association. Freezing behavior was measured as percent freezing (± SEM) across a 90-s reactivation session. ** indicates *p*<0.01.

Results of an independent-samples t-test showed no significant difference in freezing behavior between groups during the reactivation session (*t*(13) = 1.308, *p* = 0.21), suggesting that the memory in both groups was successfully retrieved (Fig 1B).

### 3.2. Destabilization of contextual fear memories is muscarinic receptor dependent

We next evaluated the involvement of muscarinic receptors in contextual fear memory destabilization using the standard procedure. A 2×2 between-subjects ANOVA revealed a significant interaction between the drug given 20 min prior to reactivation and the drug given immediately before reactivation (*F* (3,27) = 6.857, *p* = 0.004, η^2^ = 0.27). Post-hoc comparisons indicated that the VEH-MK-801 group froze significantly less than the VEH-VEH (*p* = 0.003), SCOP-VEH (*p* = 0.016), and SCOP-MK-801 (*p* = 0.004) groups during the testing session (Fig. 2A). However, when scopolamine was given 20 min prior to reactivation in conjunction with MK-801 immediately before reactivation (SCOP-MK-801), the impairing effect of MK-801 was abolished and there was no significant difference in freezing behavior compared to VEH-VEH (*p* = 0.993) and SCOP-VEH (*p* = 0.954) groups (Fig. 2A).

**Fig. 2.**
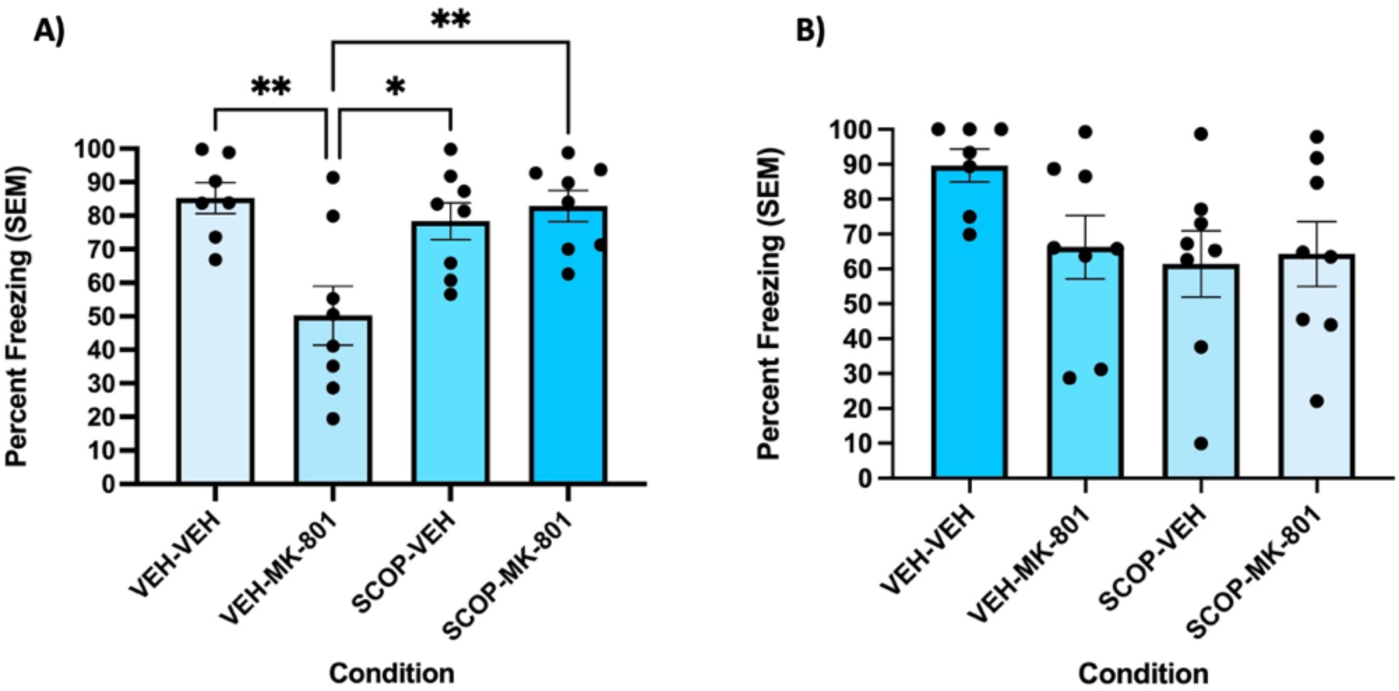
Destabilization of contextual fear memories is muscarinic receptor dependent. A) The VEH-MK-801 group froze significantly less during the test session, implying that MK-801 disrupted memory reconsolidation. SCOP-MK-801 prevented the impairing effect of MK-801. Freezing behavior (n = 8 per group) was measured as percent freezing (± SEM) across the 8-minute test session. B) The reactivation session showed no significant difference in freezing behaviour across all 4 groups. Freezing behavior was measured as percent freezing (± SEM) across the 90-s reactivation session. * indicates *p*<0.05, ** indicates *p*<0.01.

The reactivation session did not show a significant effect of drug on freezing according to a 2×2 between-subjects ANOVA (*F* (1,27) = 2.315, *p* = 0.140, η^2^ = 0.079) (Fig. 2B).

### 3.3. Modifying the fear conditioioning protocol creates a contextual fear memory that is resistant to destabilization

The purpose of this experiment was to determine whether modifying the standard fear conditoning protocol by increasing the number of shocks during training could establish a fear memory that would not readily destabilize. A 2×2 between-subjects ANOVA revealed a significant interaction between the number of shocks and the drug given (*F* (1,28) = 7.299, *p* = 0.012, η^2^ = 0.21), and Tukey’s multiple comparisons test revealed that the group receiving MK-801 and 3 shocks (3 shocks-MK-801) froze significantly less than 3 shocks-VEH (*p* = 0.015), 5 shocks-MK-801 (*p* = 0.005), and 5 shocks-VEH (*p* = 0.021) groups (Fig. 3A). However, the 5 shocks-MK-801 group did not exhibit a significant difference in freezing behavior compared to the 5 shocks-VEH (*p* = 0.943) and the 3 shocks-VEH (*p* = 0.977) groups, suggesting that the memory did not readily destabilize under standard reactivation conditions (Fig. 3A).

**Fig. 3.**
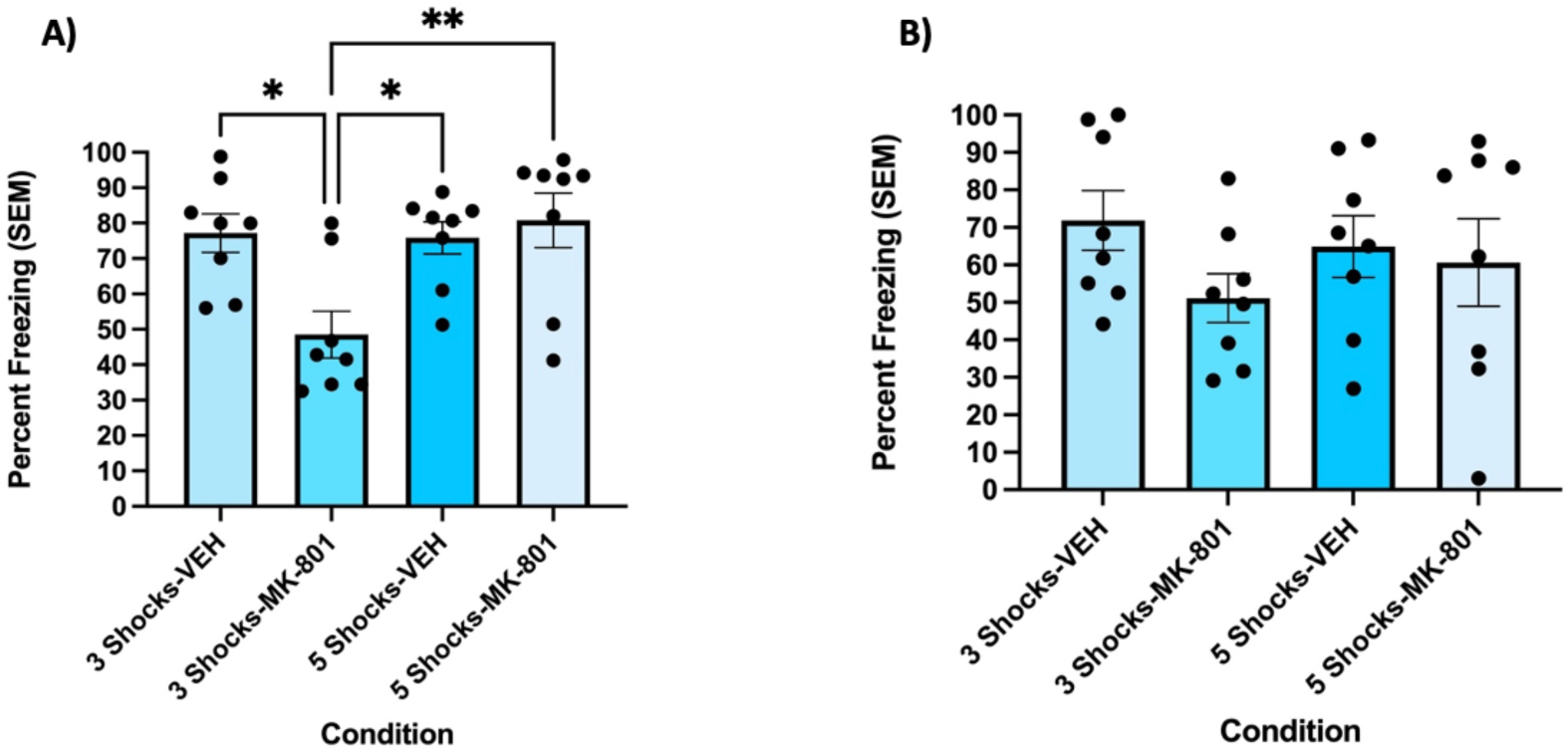
Modifying the standard fear conditioning protocol establishes a fear memory that is resistant to destabilization. A) MK-801 disrupts reconsolidation of fear memory when given to rats trained with 3 shocks, but this effect is abolished when rats are trained with 5 shocks. Freezing behavior (n = 8 per group) was measured as percent freezing (± SEM) across the 8-min test session. B) Reactivation session freezing did not indicate any significant differences between groups. Freezing behavior (n = 8 per group) was measured as percent freezing (± SEM) across the 90-s reactivation session. * indicates *p*<0.05, ** indicates *p*<0.01.

For the reactivation session, a 2×2 between-subjects ANOVA yielded no significant effect of drug (*F* (1,28) = 2.014, *p* = 0.167, η^2^ = 0.067), shock (*F* (1,28) = 0.020, *p* = 0.888, η^2^ = 0.001), or interaction between drug and shock (*F* (1,28) = 0.876, *p* = 0.357, η^2^ = 0.03) (Fig. 3B).

### 3.4. Presentation of novelty during reactivation is sufficient to induce destabilization of resistant contextual fear memory

In the previous experiment, it was shown that under standard reactivation conditions, strongly encoded fear memories are resistant to destabilization. In this experiment, we aimed to induce the destabilization of the resistant fear memory with the presentation of a novel cue during the reactivation session. Results of the 2×2 between-subjects ANOVA indicated a significant interaction between the effect of drug and the presentation of a novel cue during reactivation (*F* (1,28) = 8.591, *p* = 0.007, η^2^ = 0.24). Specifically, the group receiving a novel cue and given MK-801 immediately prior to reactivation (MK-801-NOV) froze significantly less during the test session compared to the group receiving MK-801 and no novelty (*p* = 0.038) as well as VEH and novelty (VEH-NOV) (*p* = 0.005), implying that novelty successfully promoted the destabilization of the resistant memory (Fig 4A). There was no significant difference between the MK-801 group that did not receive novelty and the VEH-NOV group (*p* = 0.846) or the VEH group that did not receive novelty (*p* = 0.964) (Fig. 4A).

**Fig. 4.**
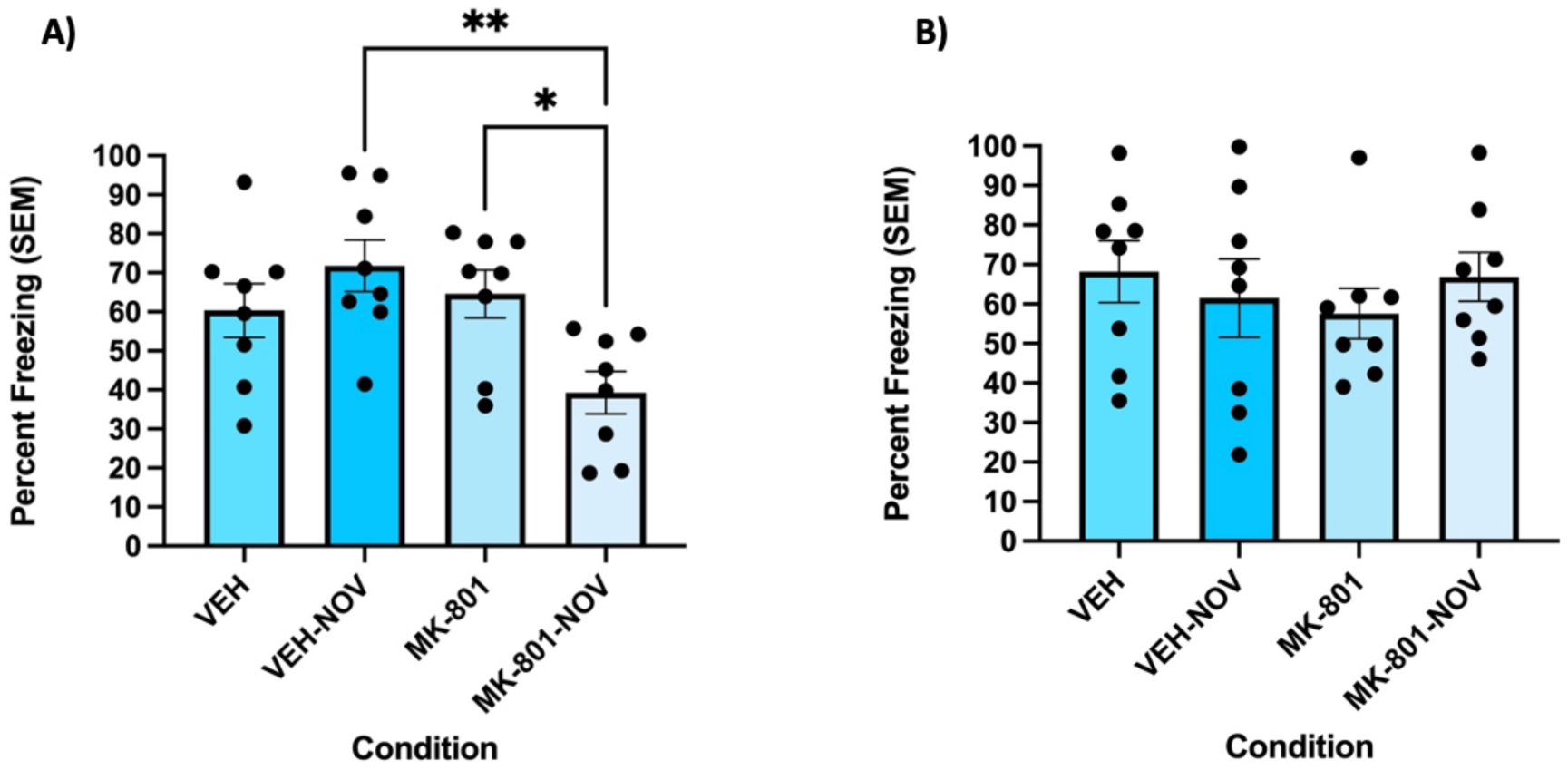
Novelty during reactivation is sufficient to promote destabilization of resistant contextual fear memory. A) The MK-801 group presented with novelty during reactivation (MK-801-NOV) froze significantly less than the group receiving MK-801 without novelty and VEH-NOV during the test session. Freezing behavior (n = 8 per group) was measured as percent freezing (± SEM) across the 8-min test session. B) No significant differences between groups were observed during the reactivation session. Freezing behavior (n = 8 per group) was measured as percent freezing (± SEM) across the 90-s reactivation session. * indicates *p*<0.05, ** indicates *p*<0.01.

There were no significant differences in freezing behaviors in the reactivation session (Fig. 4B). Results of the 2×2 between-subjects ANOVA indicated no significant interaction between presentation of novelty and drug given (*F* (1,28) = 1.072, *p* = 0.309, η^2^ = 0.037).

### 3.5. Effect of salient novelty on destabilization is blocked by mAChR antagonism

Next, we assessed whether the effect of salient novelty is dependent on mAChR activation. A 2×2 between-subjects ANOVA revealed a significant interaction between the drug given 20 minutes prior to reactivation and the drug given immediately before reactivation (*F* (1,26) = 5.736, *p* = 0.024, η^2^ = 0.18). Specifically, Tukey’s multiple comparisons test indicated that the VEH-MK-801 group froze significantly less than the VEH-VEH (*p* = 0.02), SCOP-VEH (*p* = 0.0002), and SCOP-MK-801 (*p* = 0.0032) during the testing session, which suggests that the memory destabilized when reactivated with salient novelty present and MK-801 impaired the reconsolidation of the contextual fear memory (Fig. 5A). However, when scopolamine was given 20 minutes prior to reactivation with MK-801 immediately before reactivation (SCOP-MK-801), there was no impairing effect of MK-801 (Fig. 5A).

**Fig. 5.**
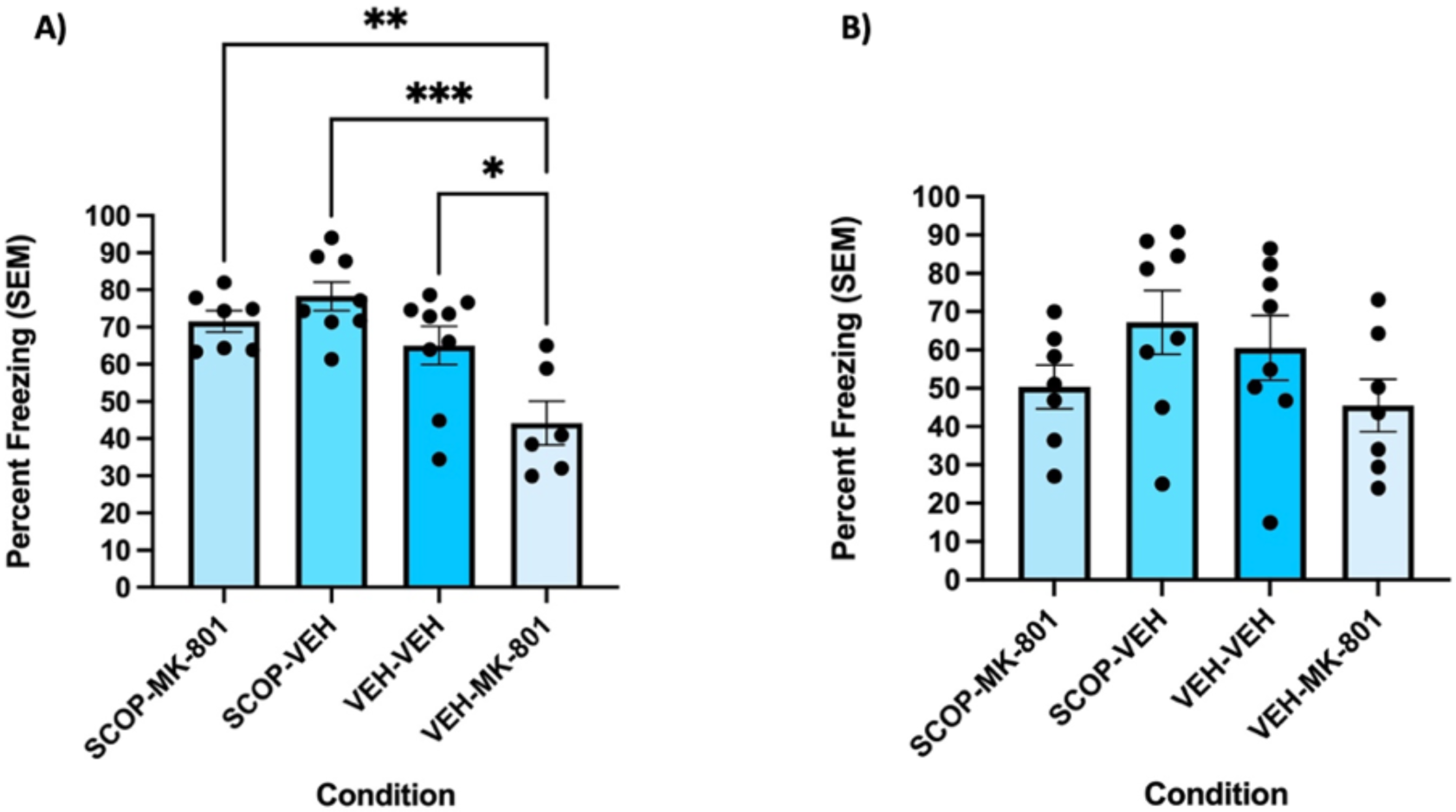
Destabilization of resistant contextual fear memories with salient novelty is muscarinic receptor dependent. A) The VEH-MK-801 group froze significantly less during the test session, implying that MK-801 disrupted memory reconsolidation. SCOP prevented the impairing effect of MK-801. Freezing behavior (n = 8 per group) was measured as percent freezing (± SEM) across the 8-min test session. B) The reactivation session indicated no significant difference in freezing behavior across all 4 groups. Freezing behavior (n = 8 per group) was measured as percent freezing (± SEM) across the 90-s reactivation session. * indicates *p*<0.05, ** indicates *p*<0.01, *** *p*<0.001.

A 2×2 between-subjects ANOVA revealed no significant interaction between the drug given 20 minutes prior and the one given immediately prior to reactivation (*F* (1,26) = 0.015, *p* = 0.905, η^2^ = 0.001) in the reactivation session (Fig 5B

### 3.6. Effect of salient novelty can be mimicked pharmacologically by M1 mAChR agonism

Finally, we were interested in studying whether activating M1 mAChRs could mimic the effect of novelty on resistant fear memory destabilization. The 2×2 between-subjects ANOVA revealed a significant interaction between the drug given 30min prior and the one given immediately prior to reactivation (*F* (1,28) = 5.292, *p* = 0.029, η^2^ = 0.159). Specifically, Tukey’s multiple comparisons test indicated that the CDD-MK-801 group froze significantly less than CDD-VEH (*p* = 0.0044), VEH-VEH (*p* = 0.0073), and VEH-MK-801 (*p* = 0.024) groups (Fig 6A). This implies that CDD promoted destabilization of strongly encoded fear memories. There was no significant freezing behavior difference between CDD-VEH and VEH-VEH (*p* = 0.997), as well as CDD-VEH and VEH-MK-801 (*p* = 0.899) (Fig 6A).

**Fig. 6.**
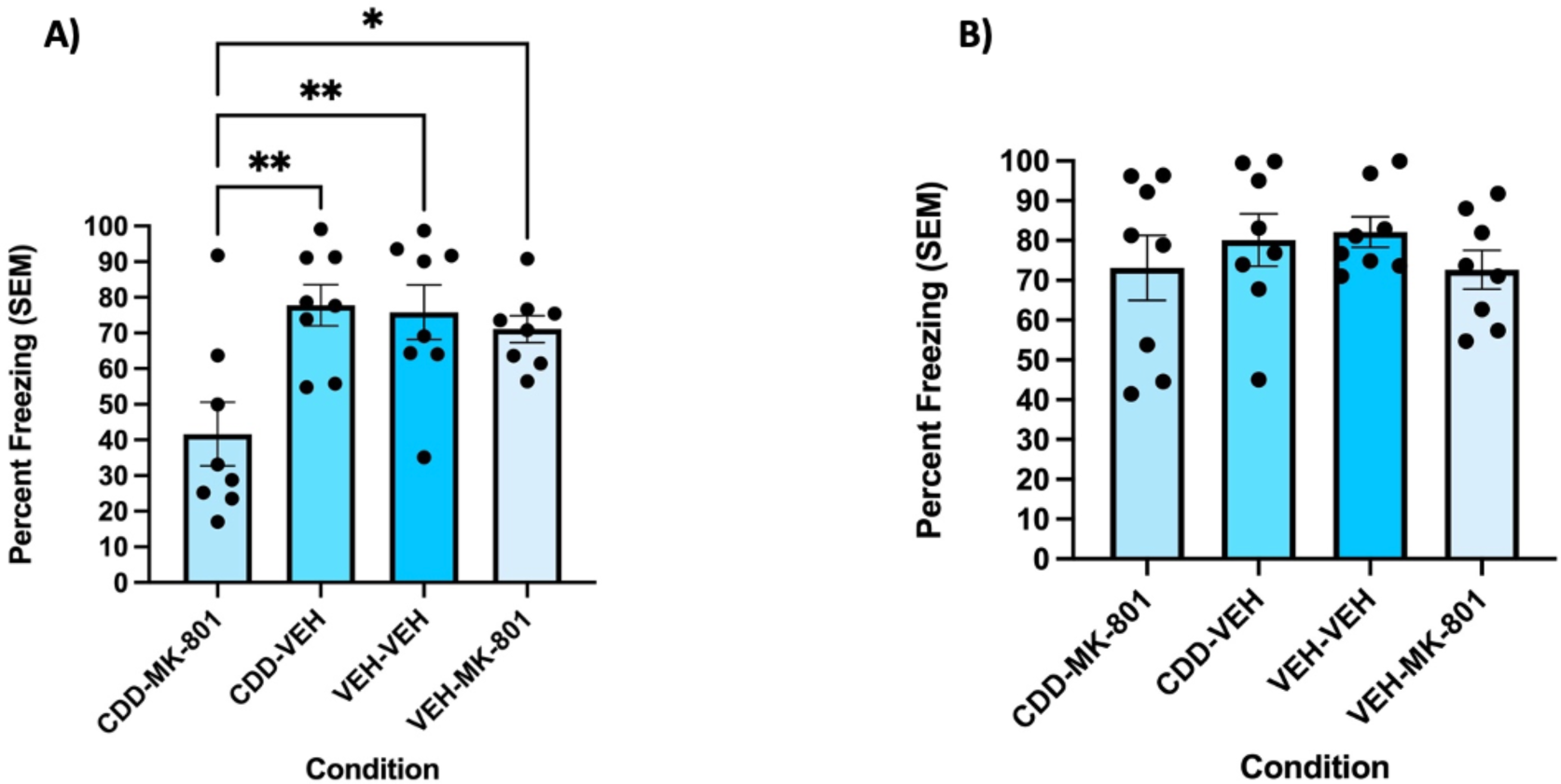
**N**ovelty-induced destabilization of resistant contextual fear memories can be mimicked using an M1 mAChR agonist. A) The CDD-MK-801 group froze significantly less during the test session, suggesting that CDD induced destabilization of the fear memory. Freezing behavior (n = 8 per group) was measured as percent freezing (± SEM) across the 8-min test session. B) The reactivation session indicated no significant difference in freezing behavior across all 4 groups. Freezing behavior (n = 8 per group) was measured as percent freezing (± SEM) across the 90-s reactivation session. * indicates *p*<0.05, ** indicates *p*<0.01.

A 2×2 between-subjects ANOVA revealed no significant interaction between the drug given 30 minutes prior and the one given immediately prior to reactivation (*F* (1,28) = 0.039, *p* = 0.845, η^2^ = 0.001) during the reactivation session (Fig 6B).

## 4. Discussion

Here, we used contextual fear conditioning to demonstrate the necessity of mAChRs in the destabilization of fear memories. First, we demonstrated that contextual fear memories readily destabilize upon reactivation when 3 context-shock pairings are used for training. To determine whether destabilization has occurred, we utilized MK-801 across all experiments to block reconsolidation. While there is evidence to support the role of NMDA receptors in destabilization and reconsolidation (Milton et al., 2013; Wideman et al., 2020), MK-801, a non-selective NMDA receptor antagonist was given immediately prior to reactivation to minimize the possibility of effects on destabilization; this procedure has been used successfully to prevent reconsolidation for various forms of memory (Brown et al., 2008; Stiver et al; Winters et al, 2009; Przybyslawski & Sara, 1997). Here, this effect was blocked, however, by pre-reactivation administration of the mAChR antagonist scopolamine, consistent with the notion that mAChR activation is necessary for contextual fear memory destabilization. As we have previously shown that mAChRs are required for the destabilization of other types of memories, such as object recognition and object location memories (Stiver et al., 2015, 2017; Huff et al., 2022; Wideman et al., 2023), this finding provides important converging evidence to support a general role for ACh in memory destabilization. To our knowledge, this is the first demonstration of the involvement of mAChRs in destabilization of fear memories.

Next, we were interested in the mechanisms underlying memory destabilization with boundary conditions. To address this, we modified our ‘standard’ CFC protocol, changing the training procedure from 3 to 5 context-shock pairings, to assess whether this change would yield a fear memory that resists destabilization. It was observed that while MK-801 again impaired reconsolidation of the relatively weak fear memory, the strongly encoded fear memory was unaffected. This suggests that the 5-shock CFC protocol established a boundary condition in which simple reactivation cues are not sufficient to induce destabilization.

Similar effects have been reported multiple times, most clearly in Wang et al. (2009) and Suzuki et al. (2004), where increasing the number of shocks prevented the amnestic drug from impairing reconsolidation. Following this demonstration, we used salient novelty during the reactivation phase to induce destabilization of the resistant fear memory, reporting that MK-801 impaired reconsolidation of the more strongly encoded fear memory under these conditions. This replicates and extends our prior findings observed in object (Stiver et al., 2015; 2017; Winters et al., 2009) and object location (Huff et al., 2022) memory tasks, where exposure to explicit contextual novelty induced destabilization of otherwise-resistant memories.

Next, we showed that scopolamine blocked novelty-induced destabilization of resistant fear memory, confirming the necessity of mAChRs in overriding the boundary condition. Lastly, we mimicked the effect of novelty using an M1 mAChR agonist, which induced destabilization of the strongly encoded contextual fear memory. We have previously shown that novelty-induced destabilization of resistant object-based memories is blocked by mAChR antagonism and mimicked by mAChR agonism (Wideman et al., 2022; Huff et al., 2022; Stiver et al., 2015; 2017), and the current findings similarly support our hypothesis that acetylcholine, acting at muscarinic cholinergic receptors, is critical for destabilization of multiple types of memories.

Here, we demonstrated that increasing the number of footshocks from three to five was sufficient to promote the formation of a contextual fear memory that was destabilization resistant. This was probably due to the increased number of training shocks establishing more prominent associative changes during encoding (Chen et al., 2021). However, the introduction of salient novelty during reactivation was sufficient to place the consolidated memory in a destabilized state. It is important to note that the novelty introduced was a tactile difference in which the floor was converted from rods to a flat surface. As such, this novel information served as a promoter for memory destabilization for modification to occur, as we have seen for object identity and object location memories in past work (Stiver et al., 2015; Huff et al., 2022). This effect is consistent with other research indicating that prediction error promotes destabilization of consolidated memories (Chen et al., 2021; Sinclair and Barense, 2018, 2019; Sinclair et al., 2021; Fernandez and Morris, 2018). Different types of prediction error, such as reactivation session time manipulation and contextual changes, have been shown to induce destabilization of different types of fear memories (Suzuki et al., 2004; Diaz-Mataix et al., 2013; Jarome et al., 2015). Yet, here we aimed to maintain the reactivation session temporal conditions and introduce a different kind of novelty as a promoter of destabilization. Importantly, it has been reported that varying levels of prediction error are required to induce destabilization of varying strengths of an established fear memory (Chen et al., 2021). In the present study, it appears that the introduction of this tactile change was sufficient to override the boundary condition introduced by two additional training shocks.

The main focus for the present study was the involvement of cholinergic transmission in the process of fear memory destabilization. We have previously shown that mAChRs, as well as nicotinic acetylcholine receptors (nAChRs), are involved in object memory destabilization (Wideman et al., 2022; Stiver et al., 2015, 2017). As such, our results build on past findings to indicate that mAChR antagonism using scopolamine blocks fear memory destabilization, consistent with a generalizable role for mAChR activation in memory destabilization and modification. It is also important to note that scopolamine did not appear to block the retrieval of fear memory in the present study. Memory impairing effects of scopolamine are typically observed when the drug is administered before learning and not when administered pre-test (Winters et al., 2006; Warburton et al., 2003). This was also addressed by Stiver et al. (2015), who showed that pre-reactivation scopolamine did not block the retrieval of object memory. Here, the drug was administered 20 min before reactivation, and subjects appeared to recall the fear memory as demonstrated by consistently intact freezing responses during the reactivation phase. Thus, blocking mAChRs appears to have specifically disrupted the destabilization process.

Acetylcholine plays a key role in memory acquisition (Hasselmo and McGaughy, 2004; Hasselmo, 2006), as well as attention and arousal (Newman et al., 2012). These established functions prompted us to hypothesize a role for ACh in novelty-induced memory destabilization of resistant contextual fear memories. The introduction of salient novel cues during reactivation should capture the attention of the rats, likely promoting the release of ACh within the hippocampus and/or amygdala, both important brain regions for contextual fear memory (Kim and Cho, 2020; Xu et al., 2016). Following the demonstration that mAChR antagonism prevented contextual fear memory destabilization, we assessed whether activating mAChRs pharmacologically could promote destabilization. Previously, we reported that infusions of CDD-0102A, an M1 mAChR agonist, in the PRh promotes the destabilization of object memories (Stiver et al., 2017). The present results similarly show that systemic CDD-0102A prior to reactivation can induce destabilization of resistant contextual fear memories. Thus, M1 mAChRs appear to play an important general role in memory destabilization. Additionally, the fact that CDD-0102A mimics the effect of salient novelty is consistent with the notion that novelty promotes ACh release and subsequent mAChR activation to induce destabilization. We have previously implicated CaMKII and ubiquitin proteasome system intracellular signaling downstream of M1 mAChR activation in object memory destabilization (Stiver et al., 2017; Wideman et al., 2023), and similar mechanisms are thought to be involved in fear memory destabilization (Lee et al 2006; Jarome and Helmstetter, 2013). Future research should address the underlying cellular mechanisms, as well as specific brain regions involved, for the mAChR-dependent fear memory destabilization demonstrated in the current study.

The present findings demonstrate an important role for cholinergic transmission in fear memory destabilization. Indeed, even resistant memories induced by a ‘strong’ training protocol can be modified when reactivated in the presence of salient novel cues and/or enhanced mAChR activation. These results are consistent with memory updating accounts of reconsolidation theory (Lee et al., 2017) and add important new insights to our understanding of the mechanistic bases of memory modification and boundary conditions. We have now demonstrated the importance of mAChR activation in destabilization and updating of object identity (Stiver et al., 2015; Winters et al., 2009), object location (Huff et al., 2022), and fear conditioned memories, strongly supporting a generalized role for cholinergic transmission in this process. The current results, in particular, if generalizable to humans, could hold important implications for potential interventions in behavioural disorders, such as post-traumatic stress disorder and phobias, characterized by pervasive, maladaptive memories (Kida, 2019).

## Authorship Contribution Statement

**Karim H. Abouelnaga:** Conceptualization, Methodology, Investigation, Formal analysis, Writing - original draft, Writing - review & editing. **Andrew E. Huff:** Investigation, Writing – original draft. **Boyer D. Winters:** Conceptualization, Methodology, Writing – review & editing, Supervision, Funding acquisition.

